# Enhanced Fear Extinction Through Infralimbic Perineuronal Net Digestion: The Modulatory Role of Adolescent Alcohol Exposure

**DOI:** 10.1101/2024.10.23.619810

**Authors:** J. Daniel Obray, Adam R. Denton, Jayda Carroll-Deaton, Kristin Marquardt, L. Judson Chandler, Michael D. Scofield

## Abstract

Perineuronal nets (PNNs) are specialized components of the extracellular matrix that play a critical role in learning and memory. In a Pavlovian fear conditioning paradigm, degradation of PNNs affects the formation and storage of fear memories. This study examined the impact of adolescent intermittent ethanol (AIE) exposure by vapor inhalation on the expression of PNNs in the adult rat prelimbic (PrL) and infralimbic (IfL) subregions of the medial prefrontal cortex. Results indicated that following AIE, the total number of PNN positive cells in the PrL cortex increased in layer II/III but did not change in layer V. Conversely, in the IfL cortex, the number of PNN positive cells decreased in layer V, with no change in layer II/III. In addition, the intensity of PNN staining was significantly altered by AIE exposure, which narrowed the distribution of signal intensity, reducing the number of high and low intensity PNNs. Given these changes in PNNs, the next experiment assessed the effects of AIE and PNN digestion on extinction of a conditioned fear memory. In Air control rats, digestion of PNNs by bilateral infusion of Chondroitinase ABC (ChABC) into the IfL cortex enhanced fear extinction and reduced contextual fear renewal. In contrast, both fear extinction learning and contextual fear renewal remained unchanged following PNN digestion in AIE exposed rats. These results highlight the sensitivity of prefrontal PNNs to adolescent alcohol exposure and suggest that ChABC-induced plasticity is reduced in the IfL cortex following AIE exposure.

## Introduction

Perineuronal nets (PNNs) are specialized components of the extracellular matrix (ECM) that are widely expressed in the brain. They consist of a hyaluronan backbone attached by link proteins to a matrix of lecticans, which are joined together by tenascin (Carulli et al., 2006; Kwok et al., 2010; Yamaguchi, 2000). Perineuronal nets play a key role in restricting plasticity and stabilizing synapses (Carulli et al., 2010; Sullivan et al., 2018). Their emergence is developmentally regulated and coincides with the closure of critical periods (Balmer et al., 2009; Cornez et al., 2020). In the medial prefrontal cortex (mPFC), PNNs initially appear early in development, and their expression continues to increase during adolescence before stabilizing in adulthood (Baker et al., 2017; Drzewiecki et al., 2020; Ueno et al., 2017). Parvalbumin interneurons (PVINs), which are crucial for initiating critical period plasticity (Fagiolini et al., 2004; Hensch et al., 1998; Takesian & Hensch, 2013), are the principal neurons enwrapped by PNNs in the cortex (Hartig et al., 1992; Lupori et al., 2023). Interestingly, the digestion of PNNs by the infusion of the enzyme Chondroitinase ABC (ChABC) into the adult brain leads to the reemergence of critical period-like plasticity (Colodete et al., 2024; Pizzorusso et al., 2002).

Perineuronal nets play a key role in regulating learning and memory (Fawcett et al., 2022; Ruzicka et al., 2022; Shi et al., 2019). In Pavlovian fear conditioning, for example, the formation of PNNs in the basolateral amygdala (BLA) prevents erasure of fear memory during extinction, while their ablation using ChABC reverses this effect (Gogolla et al., 2009; Gunduz-Cinar et al., 2019). In the hippocampus, digestion of PNNs also impacts fear learning, leading to impaired consolidation of a contextual fear memory (Hylin et al., 2013; Shi et al., 2019). Conversely, removal of PNNs in the mPFC impairs memory for cued but not contextual fear (Hylin et al., 2013). Together, these observations demonstrate that PNNs are involved in the formation and storage of fear memories.

Adolescence is a sensitive period of enhanced plasticity in the mPFC. During this period, the mPFC undergoes significant synaptic refinement (Koss et al., 2014; Petanjek et al., 2011), maturation of PVINs (Caballero et al., 2014; Cunningham et al., 2008; Du et al., 2018), and increased expression of PNNs (Baker et al., 2017; Drzewiecki et al., 2020; Ueno et al., 2017). Due to this delayed developmental profile, environmental insults, such as repeated binge-like exposure to ethanol during adolescence, can produce persistent changes in mPFC-dependent behavior and physiology. Notably, adolescent binge-like alcohol exposure has been reported to enhance fear conditioning in adult male rats (Chandler et al., 2022; Kasten et al., 2021). Given that PNNs play a key role in fear memory formation and consolidation, and considering that their expression is increased in the mPFC following AIE exposure (Dannenhoffer et al., 2022), the present study examined the impact of AIE exposure and subsequent degradation of PNNs in the infralimbic cortex on fear conditioning and extinction when tested in adulthood.

## Materials and methods

### Animals

Long-Evans dams with pups that were postnatal day (PD) 21 upon arrival were obtained from Envigo (Indianapolis, IN). On PD 23, pups were weaned and pair-housed. Rats were housed in a temperature and humidity-controlled environment on a 12 hr/12hr light/dark cycle with lights off from 09:00-21:00 each day. Food (Envigo, Teklad 2918) and water were available *ad libitum*. Only male rats were used in the present study. All procedures were approved by the Institutional Animal Care and Use Committee at the Medical University of South Carolina and carried out in accordance with the National Research Council’s Guide for the Care and Use of Laboratory Animals (National Research Council, 2011).

### Adolescent Intermittent Ethanol Exposure

Ethanol vapor exposure was performed using a standard adolescent intermittent ethanol (AIE) vapor exposure paradigm (Chandler et al., 2022; Gass et al., 2017; Landin & Chandler, 2022). Vapor exposure began on PD 28 and continued for 5 cycles until PND 44 (**Fig. 1A**). Each cycle consisted of two consecutive days of intermittent exposure with 14 hours on and 10 hours off, followed by 1-2 non-exposure days. All ethanol vapor exposure occurred in custom made chambers (Plas-Labs, Inc., Lansing, MI). Following each ethanol vapor exposure, the level of intoxication was quantified using a 5-point rating scale as follows: 1= no signs of intoxication; 2 = slightly intoxicated (slight motor impairment indicated by altered gait); 3 = moderately intoxicated (moderate motor impairment); 4 = loss of righting reflex; 5 = loss of eye blink reflex and loss of righting reflex (Gass et al., 2014; Glover et al., 2021). Ethanol vapor levels were adjusted daily based on intoxication ratings to reach a target intoxication score of 2.5 over the course of exposure. On the last day of each cycle, blood was obtained from the tail vein of each rat and analyzed for blood ethanol concentration (BEC) using an Analox AM1 alcohol analyzer (Stourbridge, GBR). After the final cycle of ethanol exposure, rats remained group housed in the vivarium until either being sacrificed at PD 78 to obtain tissue for PNN quantification (**Fig. 1A**) or undergoing surgery at approximately PD 70. Following surgery, rats were single housed until completing the fear conditioning and extinction learning paradigm at PD 94 (**Fig. 1B**).

**Figure 1.**
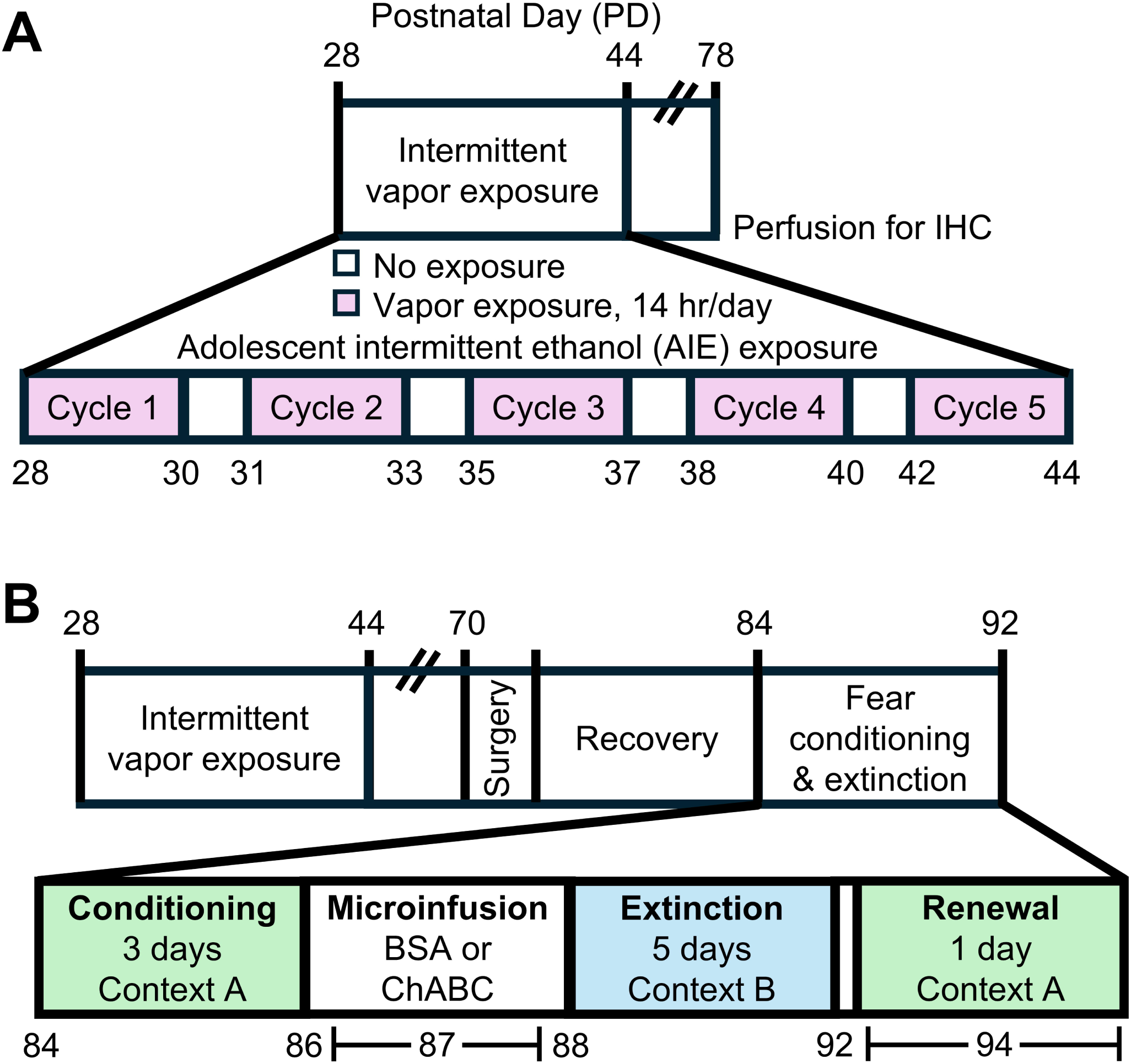
Experimental details and timeline. (**A**) Experimental timeline for perineuronal net (PNN) expression experiments. The inset image details the AIE vapor exposure paradigm. (**B**) Schema displaying the age of rats at each phase of the fear learning and extinction experiments. The inset image details the fear paradigm and timing of microinfusions.

### Quantification of Perineuronal Net Positive Cells

#### Immunohistochemistry

At PD 78, rats were deeply anesthetized using isoflurane and transcardially perfused with 4% paraformaldehyde. Brains were then removed and stored in 4% paraformaldehyde for 24 hr at 4°C before being transferred to 0.1 M phosphate buffered saline (PBS). Coronal sections (60 µm) containing the PrL/IfL were collected using a vibrating microtome (Leica Biosystems, Nussloch, DEU). Free-floating tissue sections were then blocked in 0.1 M PBS with 2% Triton X-100 (PBST) and 2% goat serum (NGS, Jackson Immuno Research, Westgrove, PA) for 2 hr with agitation. Sections were then incubated overnight with agitation at 4°C with *Wisteria floribunda* lectin (WFA, WFL, Biotinylated; Vector Laboratories, Newark, CA, B-1355-2) diluted in staining buffer (2% PBST and 2% NGS) before being washed 3 times for 5 min each in PBST. Tissue sections were next incubated for 2 hr in staining buffer containing Alexa Fluor 647, followed by three 5 min washes in PBST. Tissue sections were then mounted with ProLong Gold antifade (ThermoFisher Scientific, Waltham, MA) and coverslipped. Slides were stored at 4°C protected from light until imaging.

#### Confocal microscopy

All imaging was performed by an investigator blind to experimental conditions. A Leica TCS SP8 equipped with a Diode 638 nM laser line (Leica Microsystems, Wetzlar, DEU) was used to generate low-magnification tile scans of coronal tissue sections containing mPFC. Images were acquired using a 10X air objective (0.75 N.A.), with a frame size of 1024 X 1024 pixels, system determined step size (Nyquist-Shannon sampling), 0.75X digital zoom and a pinhole of 1 AU. Laser power and gain were first optimized and held relatively constant with minimal adjustments used only to maintain voxel saturation consistency between images.

#### Imaris Analysis

Acquired confocal images of WFA staining were imported into Bitplane Imaris (Oxford Instruments, Abingdon, GBR). At 10X magnification, WFA staining in the mPFC produces dense accumulations of labeling whose appearance is consistent with coverage of the soma and primary dendrites of putative PVINs. To quantify PNN signal at the level of individual PNNs or individual PNN positive cells, the “spots” tool was set at a diameter size of 20μm to reflect the known dimensions of typical PNN coverage of the PVIN soma and primary dendrites. Using intensity-based semi-automated deposition of 20 μm spots, PNN positive cells were counted in both the PrL and IfL cortices. Labeled PNNs were then sorted into shallow layer (layer II/III) or deep layer (layer V). Labeled PNNs within 500 µm of midline were considered layer II/III while the remainder were considered layer V. Raw counts and signal intensities of individually labeled PVIN PNNs were exported for further analysis.

### Bilateral Cannula Implantation

At approximately PD 70, a subset of rats was anesthetized with vaporized isoflurane (3%-4%) and placed in a stereotaxic alignment system (Kopf Instruments, Tujunga, CA). Bilateral microinjection guide cannula (26 GA, 1.2 mm C-C Plastics One, Roanoke, VA) were implanted to terminate 1 mm above the center of the IfL cortex at stereotaxic coordinates (from bregma): AP +3.1 mm, ML ±0.6 mm, DV −4.2 mm (Paxinos & Watson, 2014). Cannula were secured to the skull with stainless steel screws and dental cement. A removable dummy cannula (33 GA) was inserted the full length of the guide cannula. The wound was treated with topical 5% lidocaine hydrochloride and triple antibiotic ointment with bacitracin zinc, neomycin sulfate and polymyxin B sulfate. Rats were treated with a single dose of cefazolin (165 mg/kg, sq) as a prophylactic antibiotic and pre- and post-surgery with ketorolac (2.5 mg/kg, sq) for postoperative pain management. Animals were monitored daily for 7 days following surgery and began fear conditioning 1-2 weeks later.

### Fear Conditioning and Extinction Paradigm

Fear conditioning, extinction, and recall were performed using an ABA design as described previously (Chandler et al., 2022). Digitized videos were captured and analyzed by a FreezeScan Software system (Clever Sys, Reston, VA). Briefly, on each of 3 days, conditioning consisted of a 60 s acclimation period followed by four tones (conditioned stimulus [CS], 2 kHz, 80 dB, 30 s) co-terminating with a 2 s foot shock (unconditioned stimulus [US], 0.75 mA). Each tone-shock pairing was separated by a 10 s interval. In a visually distinct context, beginning 48 hours after the final day of conditioning, rats underwent a 5-day extinction protocol. Each extinction session consisted of a 60 s acclimation period followed by 10 CS presentations with no co-terminating foot shock. Each CS presentation was separated by a 10 s inter-stimulus interval. Forty-eight hr after the last extinction session, rats were returned to the conditioning context and tested for context-dependent fear renewal using a single CS presentation.

### Chondroitinase ABC infusion

One day after the final conditioning session and 24 hr prior to the first day of extinction, rats were microinfused with either Chondroitinase ABC (ChABC, 0.18 U/µl, 100 nl, Sigma-Aldrich, Burlington, MA, C3667) reconstituted in 0.01% bovine serum albumin (BSA) as per manufacturer’s instructions, or BSA vehicle control (0.01%,100 nl, Sigma-Aldrich, A9647). This dose of ChABC was used to assure complete digestion of PNNs based on previously published findings (Steullet et al., 2014; Vo et al., 2013). After lightly restraining the rat, the dummy cannula was removed and a sterile 33-gauge internal cannula (Plastics One) that extended 1 mm past the end of the implanted guide cannula inserted. The internal infusion cannula was connected to a 1 µL Hamilton syringe (Hamilton Company, Reno, NV, 80100) driven by a microinfusion pump (Harvard Apparatus, Holliston, MA). Chondroitinase ABC or BSA vehicle was delivered at a rate of 1 nl/s, for 100 s to deliver a total of 100 nl. The infusion cannula was left in place for 2 min after infusion to allow for solution diffusion.

### Histological Verification of Chondroitinase ABC Lesion

Twenty-four hr after the completion of the behavioral protocol, rats were deeply anaesthetized with isoflurane and transcardially perfused with 4% paraformaldehyde. Brains were removed, immersed in 4% paraformaldehyde for 24 hr and then transferred to a 30% (w/v) sucrose solution for at least 72 hr at 4 °C. Brains were flash frozen and stored at −80 °C for at least 24 hr prior to slicing into 40 mm coronal sections on a cryostat (Thermo Fisher Scientific, Waltham, MA, Microm HM 550). Successful degradation of PNNs was determined by colorimetric 3-3’-Diaminobenzidine staining with recognition by biotinylated WFA (Vector Laboratories) through use of the VECTASTAIN Elite ABC-HRP avidin-biotin complex (Vector Labs, PK6200). Eight rats were excluded from the study because the ChABC lesion spread significantly beyond the IfL.

### Statistical Analysis

A total of 10 rats (6 tile images per animal) were used for the PNN expression studies. For analysis of individual PNN intensities, absolute frequency distributions were generated from exported PNN intensity data. Using GraphPad Prism 10.3 (GraphPad Software, Boston, MA), a chi-square test of independence was performed to determine distributional shifts in intensity distributions. When significant, the chi-square test was followed by Bonferroni corrected two-sample tests of proportion, performed at each intensity bin to characterize the AIE-induced shifts in the frequency distribution. For analysis of PNN count data, nested ANOVAs were conducted using Stata 15.1 (StataCorp, College Station, TX) with each region and layer analyzed separately. Freezing data from the conditioning and extinction phases were analyzed using separate mixed model ANOVAs. Freezing data from the renewal session was analyzed using a two-way ANOVA. The Greenhouse-Geisser correction for sphericity was applied to all repeated measures factors. For post-hoc multiple comparison tests, the Bonferroni method was used to control the family-wise error rate. A *p*-value of < 0.05 was considered the threshold for statistical significance.

## Results

This study utilized a well-validated vapor inhalation model of ethanol exposure that mimics repeated episodes of binge-like ethanol exposure (Gass et al., 2014; Glover et al., 2021). Behavioral intoxication and blood ethanol concentration (BEC) were assessed at the end of each exposure cycle. The average behavioral intoxication score using the 5-point rating scale was 2.3 ± 0.2, which represents a moderate level of intoxication. The coinciding BEC value was 206.6 ± 13.0 mg/dl.

The impact of AIE exposure on the number of PNNs in the PrL and IfL subregions of the male rat prefrontal cortex was examined using WFA staining. Representative confocal images of PNN staining are presented in **Figure 2**. Analysis of PNN staining, and quantification of PNN positive cells in the PrL subregion revealed an increased number of PNN positive cells in AIE-exposed rats compared to Air control rats (*F*_(1,8)_ = 8.97, *p* = 0.0172, partial η^2^ = 0.5286 [0.0189, 0.7425]; **Fig. 3A**). When subdividing the analysis by cortical layer, we found a significant AIE-induced increase in PNN positive cells in layer II/III and a trend toward an increase in layer V (layer II/III: *F*_(1,8)_ = 18.75, *p* = 0.0025, partial η^2^ = 0.7010 [0.1657, 0.8357]; layer V: *F*_(1,8)_ = 3.91, *p* = 0.0834; **Fig. 3B**). In contrast, evaluation of the IfL subregion revealed no differences in the number of PNN positive cells between AIE exposed and Air control rats (*F*_(1,8)_ = 1.60, *p* = 0.2415; **Fig. 3C**). While layer-specific analysis revealed no change in number of PNN positive cells in IfL layer II/III, there was a significant decrease in layer V in AIE-exposed rats (layer II/III: *F*_(1,8)_ = 1.36, *p* = 0.2778; layer V: *F*_(1,8)_ = 8.84, *p* = 0.0178, partial η^2^ = 0.5250 [0.0172, 0.7405]; **Fig. 3D**).

**Figure 2.**
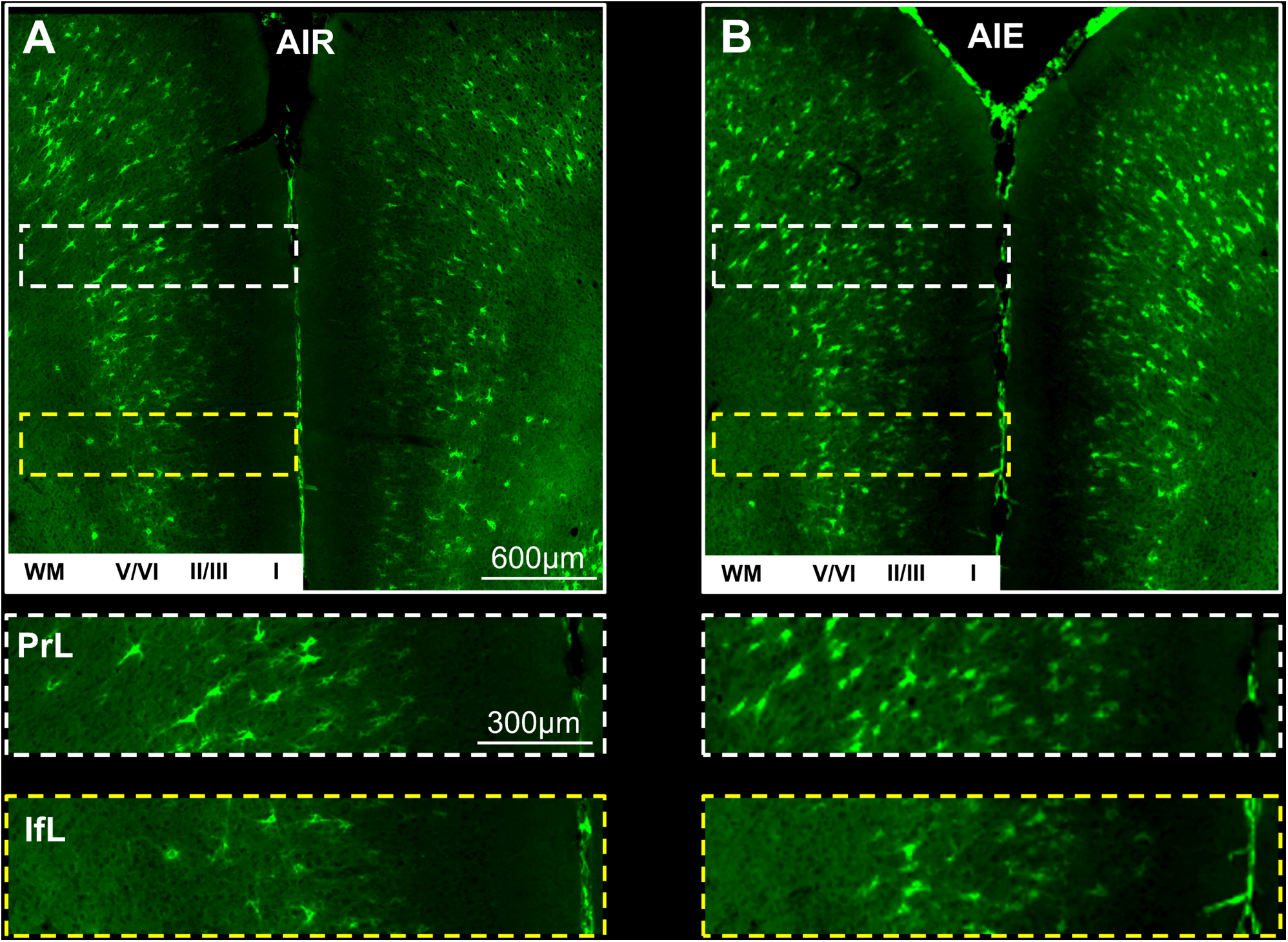
Representative Images of cortical perineuronal net staining in air and adolescent intermittent ethanol exposed animals. **(A)** Representative confocal images of Air (control) animals. A section of tissue within one hemisphere in the prelimbic (PrL) cortex is highlighted with a white hashed rectangle. A section of infralimbic (IfL) cortex is highlighted with a yellow hashed rectangle. Also included is an indication of cortical layer in a white box. Below the 10X magnification image depicting both hemispheres, inset panels are shown representing a single hemisphere of each cortical subregion, shown at higher magnification. The inset panels include corresponding white (PrL) and yellow (IfL) hashed outlines **(B)** Representative confocal images of AIE animals are organized and depicted as in **(A)**.

**Figure 3.**
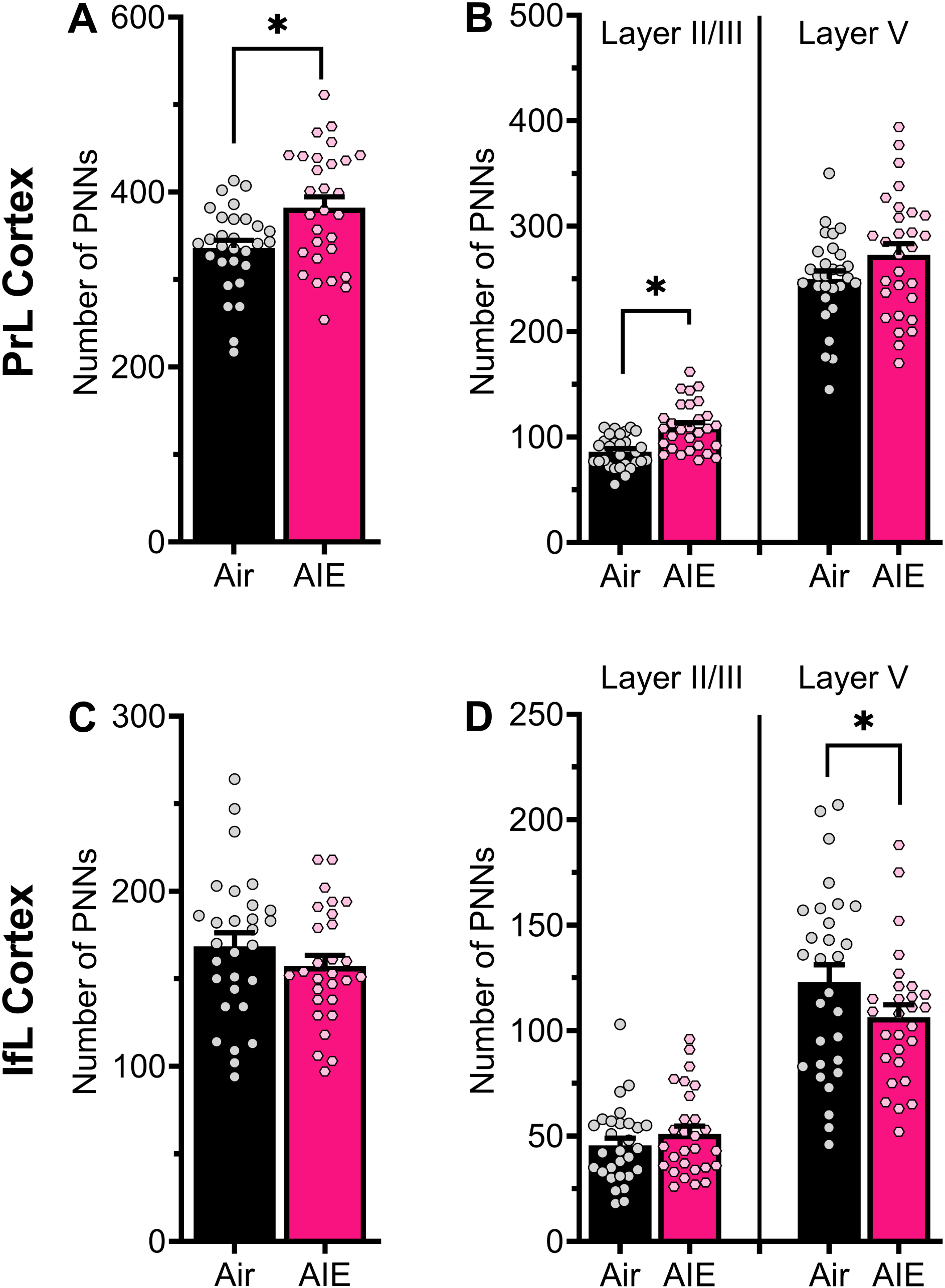
Adolescent intermittent ethanol exposure bidirectionally modulated perineuronal net numbers in the prelimbic and infralimbic cortex. (**A**) Adult male rats exposed to adolescent intermittent ethanol (AIE) displayed increased numbers of perineuronal net (PNN)-positive neurons in the prelimbic (PrL) cortex. (**B**) Layer-specific analyses revealed that AIE-exposure led to a significant increase in PNN numbers in layer II/III, while layer V showed a non-significant trend toward increased PNN numbers. (**C**) In the infralimbic (IfL) cortex, there was no significant difference in PNN counts between AIE exposed rats and Air control rats. (**D**) Within the IfL, AIE did not significantly affect the number of PNN-positive cells in layer II/III, however, in layer V AIE decreased the number of these cells. * denotes *p* < 0.05. n = 5 rats/group with 5-6 replicates/animal.

The next set of analyses assessed the effect of AIE exposure on the staining intensity of PNNs at the level of individual cells. Analysis of PNN positive cells in the PrL cortex revealed that AIE exposure significantly altered the frequency distribution of PNN staining intensity (Χ^2^_(15)_ = 243.3519, *p* < 0.001, *V* = 0.1081; **Fig. 4A-B**). In the IfL cortex, a similar AIE-induced change in the distribution of PNN intensity was observed (Χ^2^_(14)_ = 218.1251, *p* < 0.001, *V* = 0.1600; **Fig. 4C-D**). In both regions, the alteration in the intensity frequency distribution reflected a reduction in the proportion of the strongest and weakest stained PNNs. Consistent with this, the spread of PNN staining intensity was significantly reduced in both the PrL (Brown-Forsythe test: *F*_(1,20827)_ = 176.9881, *p* < 0.0001) and the IfL (Brown-Forsythe test: *F*_(1,8523)_ = 195.2209, *p* < 0.0001) of AIE exposed rats.

**Figure 4.**
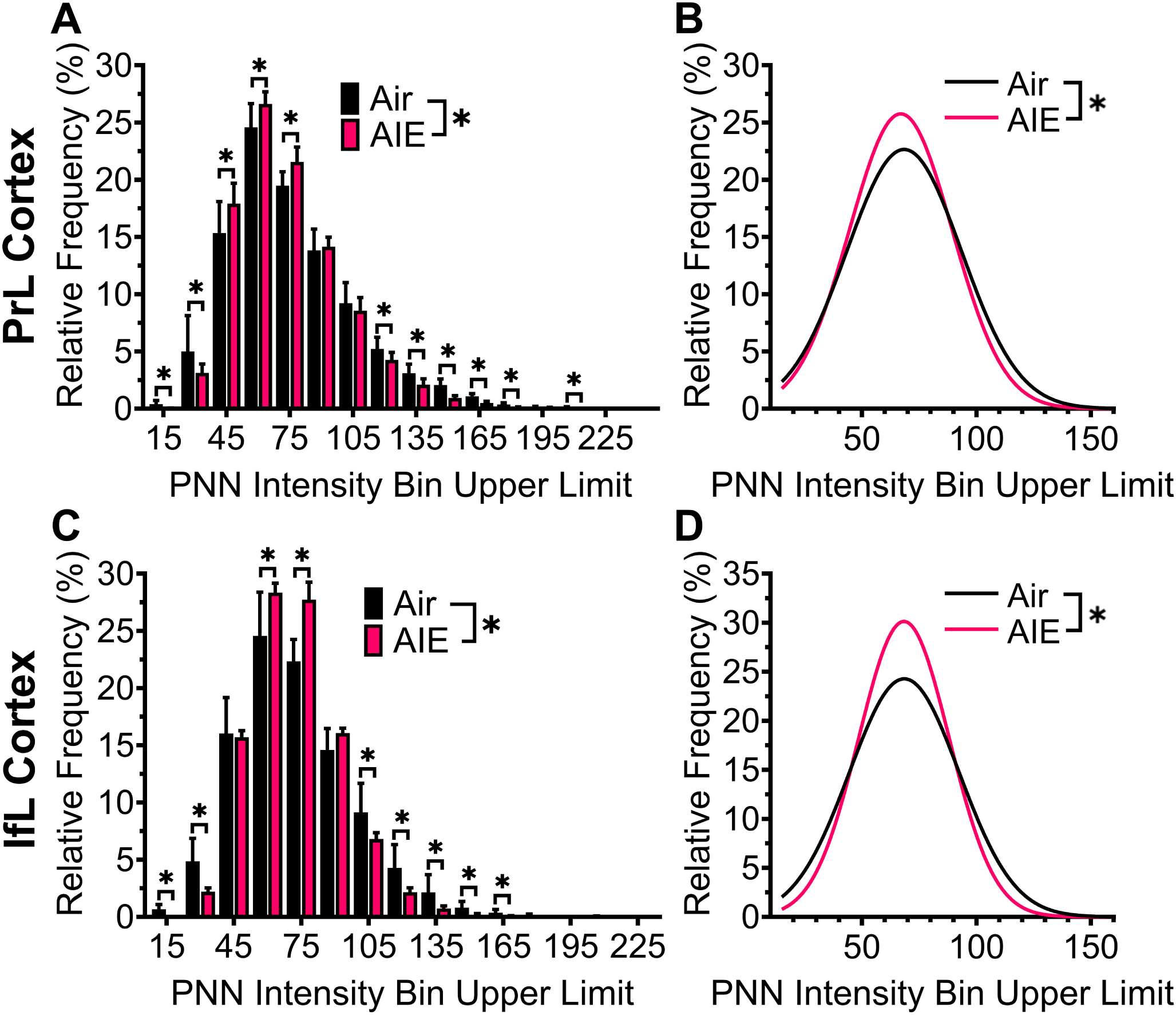
Adolescent intermittent ethanol exposure shifted the distribution of perineuronal net staining intensity in the prelimbic and infralimbic cortex. (**A**) Relative frequency of *Wisteria floribunda* agglutinin (WFA) staining intensity of individual perineuronal nets (PNNs) in the prelimbic (PrL) cortex. Within the PrL cortex, adolescent intermittent ethanol (AIE) exposure increased the relative frequency of intermediate intensity PNNs. (**B**) Gaussian curves showing the decreased variability of PNN intensity and the increased frequency of intermediate intensity PNNs in the PrL cortex of AIE exposed rats. (**C**) Relative frequency of individual PNN staining intensity in the infralimbic (IfL) cortex. Within the IfL cortex, the distribution of PNN staining intensity was altered by AIE exposure, reflecting an AIE-induced increase in the proportion of intermediate intensity PNNs. (**D**) Gaussian curves displaying the decreased variability of PNN intensity and the increased frequency of intermediate intensity PNNs in the IfL cortex of AIE exposed rats. * denotes *p* < 0.05. For the PrL, n = 9,745 PNNs (Air) and 11,084 PNNs (AIE). For the IfL, n = 4,509 PNNs (Air) and 4,016 PNNs (AIE). There were 5 rats/group.

It is well documented that PNNs have a critical role in plasticity including learning and memory, with numerous studies demonstrating their involvement in both fear conditioning and extinction learning (Banerjee et al., 2017; Gogolla et al., 2009; Gunduz-Cinar et al., 2019; Liu et al., 2023; Poli et al., 2023; Shi et al., 2019; Thompson et al., 2018). Additionally, the IfL cortex is recognized as a key brain region for extinction learning (Bukalo et al., 2015; Do-Monte et al., 2015; Morgan & LeDoux, 1995; Morgan et al., 1993). Therefore, the next set of studies investigated the impact of digestion of PNNs in the IfL cortex on extinction of a conditioned fear memory. A set of pilot experiments were first conducted to qualitatively assess the time-course of PNN depletion and recovery following microinfusion of ChABC into the IfL cortex. This confirmed that microinfusion of ChABC into the IfL cortex led to the rapid and nearly complete degradation of PNNs within the boundary of the injection site, which was clearly observable 24 hr post-infusion (**Fig. 5A**). Consistent with previous reports (Bruckner et al., 1998; Lensjø et al., 2017), PNN staining gradually reemerged beginning by 7 days post-infusion and continuing through 21 days (**Fig. 5A**).

**Figure 5.**
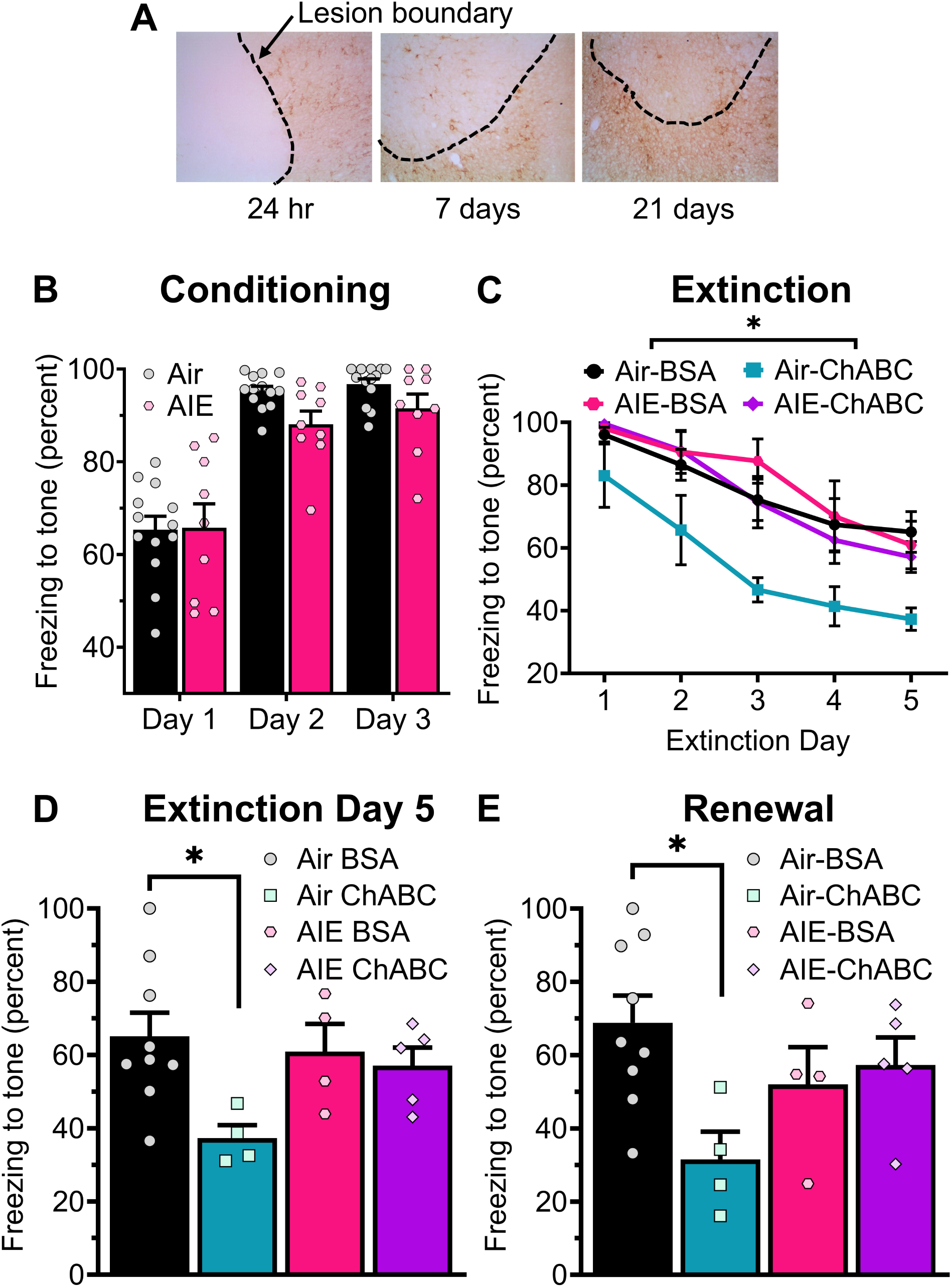
Depletion of infralimbic perineuronal nets enhanced extinction learning: modulation by adolescent alcohol exposure. (**A**) Images depicting *Wisteria floribunda* agglutinin (WFA) staining of perineuronal nets 24 hr, 7 days, and 21 days after microinfusion of chondroitinase ABC (ChABC). Significant digestion of PNNs is evident 24 hr after the microinfusion. At 7 days post-infusion limited recovery of PNNs is apparent, with a more complete recovery of PNN levels visible at 21 days post-infusion. (**B**) Depiction of the increase in freezing to the tone that occurred across three fear conditioning sessions in which the tone was paired with a foot shock. There were no significant differences between Air control and adolescent intermittent ethanol (AIE) exposed rats in acquisition of the tone-shock pairing. (**C**) Graphic displaying the gradual decline in freezing to the tone that occurred over 5 extinction learning sessions in which the tone was presented with an accompanying foot shock. Extinction learning was significantly enhanced in rats that received a microinfusion of ChABC into the infralimbic cortex. (**D**) Graphic representation of freezing in response to the tone during the final conditioned fear extinction session. Adolescent alcohol exposure prevented the enhanced extinction learning that occurred in Air control rats following the digestion of PNNs in the IfL cortex. (**E**) Depiction of freezing in response to a single tone presentation during the context-dependent fear renewal task. Digestion of PNNs by ChABC significantly reduced fear renewal in Air control but not AIE exposed rats. * denotes *p* < 0.05. n = 9 (Air + bovine serum albumin [BSA]), 4 (Air + ChABC), 4 (AIE + BSA), 5 (AIE + ChABC).

After verifying that PNNs remained substantially depleted within a 7 day window following ChABC infusion, the next set of studies examined the impact of PNN digestion on extinction of a conditioned fear memory in Air control and AIE-exposed rats. For these experiments, both Air control and AIE-exposed rats underwent a 3 day tone-shock fear conditioning procedure.

Analysis of the fear conditioning sessions confirmed that the rats learned the tone-shock association, as indicated by an increase in tone-induced freezing across sessions (main effect session: *F*_(2,36)_ = 89.34, *p* < 0.0001, partial η^2^ = 0.8323 [0.7045, 0.8805]; **Fig. 5B**). There were no significant differences in the acquisition of the tone-shock pairing based on AIE exposure, treatment group (BSA vs ChABC), or any interaction between AIE, treatment group, and session (main effect AIE: *F*_(1,18)_ = 1.34, *p* = 0.2624; main effect ChABC: *F*_(1,18)_ = 0.14, *p* = 0.7171; session x AIE interaction: *F*_(2,36)_ = 2.03, *p* = 0.1462; session x ChABC interaction: *F*_(2,36)_ = 1.60, *p* = 0.2162; session x AIE x ChABC interaction: *F*_(2,36)_ = 0.12, *p* = 0.8858).

On the day following the final fear conditioning session, rats received a bilateral infusion of either vehicle (saline-BSA) or ChABC. Beginning 24 hr after the infusion, the rats then underwent daily extinction sessions in a novel context for 5 consecutive days. During each session, the tone was presented 10 times without the previously paired foot-shock and freezing in response to the tone was measured. Analysis of these data confirmed that extinction learning took place, as tone-induced freezing significantly decreased across sessions (main effect session: *F*_(4,72)_ = 51.10, *p* < 0.0001, partial η^2^ = 0.7395 [0.6155, 0.7941]; **Fig. 5C**). Consistent with previous reports on enhanced plasticity associated with degradation of PNNs, the rate of fear extinction was augmented by PNN digestion (main effect ChABC: *F*_(1.18)_ = 5.27, *p* < 0.0339, partial η^2^ = 0.2266 [0.0000, 0.4879]). However, there was no significant effect of AIE exposure, or any interaction between session, ChABC, and AIE on extinction learning (main effect AIE: *F*_(1.18)_ = 4.27, *p* = 0.0536; ChABC x AIE interaction: *F*_(1.18)_ = 2.35, *p* = 0.1425; session x ChABC interaction: *F*_(4,72)_ = 1.72, *p* = 0.1950; session x AIE interaction: *F*_(4,72)_ = 1.18, *p* = 0.3187; session x ChABC x AIE interaction: *F*_(4,72)_ = 0.19, *p* = 0.8216). A post-hoc analysis of freezing in response to the tone on the fifth extinction day further revealed that PNN digestion reduced freezing in Air control (*t*_(18)_ = −2.63, *p* = 0.034, Hedges’s *g* = 1.5196 [0.2303, 2.7572]) but not AIE exposed rats (*t*_(18)_ = −0.32, *p* = 1.000). Therefore, rats that had been exposed to AIE failed to exhibit facilitation of fear extinction associated with digestion of PNNs in the IfL cortex.

Two days after the final extinction session, context-dependent renewal of the conditioned fear memory was assessed by measuring freezing behavior when the animals were placed back in the original fear conditioning context. Analysis revealed a significant ChABC x AIE interaction (ChABC x AIE interaction: *F*_(1.18)_ = 5.71, *p* < 0.0281, partial η^2^ = 0.2407 [0.0000, 0.4995]; **Fig. 5D**), but no significant effects of ChABC or AIE exposure alone (main effect ChABC: *F*_(1.18)_ = 3.22, *p* < 0.0895; main effect AIE: *F*_(1.18)_ = 0.25, *p* < 0.6218). Similar to what was observed with extinction learning, post-hoc analysis further indicated that ChABC reduced fear renewal in Air control rats (*t*_(18)_ = −3.14, *p* = 0.011, Hedges’s *g* = 1.681 [0.358, 2.951]) but had no effect in AIE exposed rats (*t*_(18)_ = 0.40, *p* = 1.000).

## Discussion

Repeated binge-like exposure to alcohol during adolescence is associated with increased PNN density in the mPFC of adult male rats (Dannenhoffer et al., 2022). This study investigated how AIE exposure affects numbers of PNN positive cells and WFA stain intensity in layer II/III and V of the PrL and IfL subregions of the mPFC. As PNNs play a key role in learning and memory, the study also examined the impact of AIE exposure combined with ChABC digestion of IfL PNNs on fear memory and extinction learning. Key findings from these experiments include: AIE exposure increased the number of PNN-positive cells in the PrL but decreased them in layer V of the IfL; AIE altered the frequency distribution of PNN staining intensity in both the PrL and IfL; and AIE exposure prevented PNN digestion in the IfL cortex from enhancing extinction learning.

In the PrL, there was a significant increase in the number of PNN positive cells in layer II/III, with a non-significant trend towards an increase in layer V. In contrast, the number of PNN positive cells remained unchanged in layer II/III of the IfL, while they decreased in layer V due to AIE exposure. Previous studies examining the effects of adolescent alcohol exposure on PNN expression in the mPFC have yielded mixed results, with one study reporting enhanced PNN density in both the IfL and PrL of male rats (Dannenhoffer et al., 2022), but another reporting no changes in either male or female rats (Sullivan et al., 2024). Notably, in the latter study by Sullivan and colleagues, the rats underwent behavioral training prior to evaluating the impact of AIE exposure on the density of PNN positive neurons. Given that PNNs are subject to experience-dependent remodeling (Banerjee et al., 2017; Dankovich et al., 2021; Van den Oever et al., 2010), this may help explain the discrepancies between these studies. In addition to quantifying the effects of adolescent alcohol exposure on number of PNN positive cells, the present study also assessed its effect of on the staining intensity of PNN positive cells. In both the PrL and IfL cortex, AIE exposure narrowed the intensity frequency distribution of PNN staining by reducing the proportions of the most and least intensely stained cells while increasing the frequency of cells in the mid-range of staining intensity. These findings underscore the nuanced effects of adolescent alcohol exposure on PNNs in the rat mPFC.

The mechanism by which AIE alters PNN expression is not known. Although speculative, a potential explanation is that it may be related to disruption of dopamine 1 (D1) receptor dependent remodeling of the ECM. Activation of D1 receptors has been shown to stimulate activity of matrix metalloproteinase (MMP) and a disintegrin and metalloproteinase with thrombospondin motifs (ADAMTS) 4 and 5, which contribute to PNN remodeling (Li et al., 2016; Mitlöhner et al., 2020). Given that adolescent alcohol exposure reduces D1 modulation of neuronal excitability (Obray et al., 2022; Trantham-Davidson et al., 2017), it may also decrease D1 stimulation of MMPs and ADAMTS4 and 5, leading to more static PNNs due to reduced proteolysis of PNN components. Another possibility is that AIE alters the activity of receptor protein tyrosine phosphatase (RPTP) ζ in the mPFC. Receptor protein tyrosine phosphate ζ and its splice variant phosphacan, are crucial for anchoring PNNs to the cell surface through interactions with tenascin-R (Eill et al., 2020). Notably, RPTP ζ regulates PNN intensity (Chu et al., 2018; Galan-Llario et al., 2023), and inhibiting RPTP ζ blocks AIE-induced reductions in the most intensely stained hippocampal CA1 PNNs (Galan-Llario et al., 2024), suggesting a potential role for RPTP ζ in the AIE-induced changes observed in PNNs.

Fear conditioning and extinction involve multiple brain regions, including the mPFC. Within the mPFC, the PrL and IfL subregions modulate fear expression bidirectionally through their projections to the BLA and thalamus (Marek et al., 2019; Sung & Kaang, 2022). Perineuronal nets are important for learning and memory, and their digestion in the mPFC disrupts fear recall (Hylin et al., 2013). However, the role of prefrontal PNNs in fear extinction has not been extensively studied. The present study examined the effects of AIE exposure, in conjunction with ChABC digestion of IfL PNNs, on conditioned fear extinction learning and context-dependent renewal. These experiments revealed that neither fear conditioning nor extinction learning were significantly impacted by adolescent alcohol exposure. However, digestion of PNNs in the IfL cortex by infusion of ChABC was found to enhance extinction learning and reduce context-dependent fear renewal. Overall, these findings align with previous studies on AIE and fear, which found no significant effect of adolescent alcohol exposure on extinction learning, along with either no change (Bergstrom et al., 2006; Broadwater & Spear, 2013) or enhanced fear conditioning (Chandler et al., 2022; Kasten et al., 2021). Notably, while previous research indicated that AIE exposure attenuated context renewal (Chandler et al., 2022), the present study found no significant effect of AIE on renewal in BSA (vehicle) treated rats. There are methodological differences between studies – such as testing extinction recall prior to evaluating renewal and the microinfusion of BSA – which may contribute to the absence of a significant effect of AIE on renewal.

Fear extinction is a plasticity dependent process. During fear extinction, activation of pyramidal neurons in the IfL enhances extinction learning and subsequent recall (Do-Monte et al., 2015; Thompson et al., 2010), whereas activation of IfL PVINs impairs extinction recall (Binette et al., 2023). Perineuronal nets enwrap cortical PVINs (Hartig et al., 1992; Lupori et al., 2023) and play a key role in regulating their physiology (Chu et al., 2018; Stevens et al., 2021; Tewari et al., 2024). Loss of PNNs reduces the intrinsic excitability of PVINs and both decreases the amplitude and increases the frequency of spontaneous excitatory postsynaptic currents (sEPSCs) at PVINs (Gonzalez et al., 2022). These changes lead to disinhibition of pyramidal neurons (Slaker et al., 2015). Furthermore, extinction learning induces neuregulin 1 (NRG1) ErB-B2 receptor tyrosine kinase 4 (ErbB4) dependent plasticity in IfL PVINs, which disinhibits IfL pyramidal neurons (Chen et al., 2022). Thus, the observed effect of PNN degradation may be linked to enhanced activation of IfL pyramidal neurons during extinction trials. Interestingly, RPTP ζ suppresses NRG1-ErbB4 signaling (Buxbaum et al., 2008; Fujikawa et al., 2007; Takahashi et al., 2011; Tanga et al., 2019), suggesting that PNN digestion might also enhance extinction learning by augmenting NRG1-ErbB4 signaling. However, this remains speculative, especially since PNNs are necessary for NRG1-ErbB4 mediated depression of inhibitory neurotransmission in the hippocampus (Dominguez et al., 2019). Another interesting observation in the present study was that AIE blocked facilitation of fear extinction and renewal following digestion of PNNs in the IfL cortex. It is interesting to speculate that AIE exposure may have negatively impacted plasticity processes that are engaged by PNN digestion in the IfL cortex.

One of the limitations of the present study is that it did not include female rats. Recent research has highlighted sex differences in the effects of AIE on conditioned fear (Chandler et al., 2022; Kasten et al., 2020; Kasten et al., 2021; Landin & Chandler, 2022) and has identified behavioral variations between male and female rats in their conditioned fear responses (Gruene et al., 2015; Mitchell et al., 2022). Future follow-up studies should include parameters optimized for female rats to better assess the interactions between AIE and PNNs on extinction learning. Another caveat of the study is the use of ChABC to digest PNNs. While ChABC effectively degrades PNNs, it is not specific for them, as the components it degrades are also found in other parts of the ECM. Consequently, the effects of ChABC may arise from its actions on both PNNs and the broader ECM.

## Conclusion

The first set of experiments in this study examined the effect of AIE exposure on the number of PNN-positive cells in the PrL and IfL subregions of the mPFC. In the PrL cortex, it was observed that AIE increased the overall number of PNN-positive cells. This effect was layer specific, with a significant increase in the number of PNN-positive cells in Layer II/III but no change in Layer V. In the IfL cortex, while AIE did not change the overall density of PNN-positive cells, it did produce a significant reduction in the number of PNN-positive cells in Layer V but no change in Layer II/III. Further analysis of the staining intensity for PNN positive cells revealed a narrowed distribution whereby AIE increased the proportion those cells with intermediate PNN staining intensity in both the PrL and IfL. The second set of experiments in this study assessed the combined effects of AIE and ChABC digestion of IfL PNNs on fear extinction and renewal. Notably, digestion of PNNs in the IfL cortex enhanced conditioned fear extinction learning and reduced context-dependent fear recall. However, both effects were blunted by AIE exposure. These findings contribute to a growing body of research indicating that PNN expression is sensitive to adolescent alcohol exposure. Additionally, the results suggest that AIE exposure may negatively impact the plasticity triggered by PNN digestion in the IfL cortex.

## Acknowledgements

We would like to thank members of the Chandler lab for their help in conducting the ethanol vapor exposure. This work was supported by National Institute of Health grants AA027706 (LJC), AA019967 (LJC), and AA027427 (KM). The authors declare that the research was carried out in the absence of any commercial or financial relationships that could be construed as a conflict of interest.

## CRediT Authorship Contribution Statement

**J. Daniel Obray**: Formal analysis, Visualization, Writing – original draft, and Writing – review and editing. **Adam R. Denton**: Formal analysis, Visualization, Writing – original draft, and Writing – review and editing. **Jayda Carroll-Deaton**: Investigation. **Kristin Marquardt**: Conceptualization, Formal Analysis, and Investigation. **L. Judson Chandler**: Conceptualization, Funding acquisition, Supervision, Visualization, Writing – original draft, and Writing – review and editing. **Michael D. Scofield**: Conceptualization, Supervision, Visualization, Writing – original draft, and Writing – review and editing.

